# UFBoot2: Improving the Ultrafast Bootstrap Approximation

**DOI:** 10.1101/153916

**Authors:** Diep Thi Hoang, Olga Chernomor, Arndt von Haeseler, Bui Quang Minh, Le Sy Vinh

**Author notes:** Shared first authors (Diep Thi Hoang, Olga Chernomor).

## Abstract

The standard bootstrap (SBS), despite being computationally intensive, is widely used in maximum likelihood phylogenetic analyses. We recently proposed the ultrafast bootstrap approximation (UFBoot) to reduce computing time while achieving more unbiased branch supports than SBS under mild model violations. UFBoot has been steadily adopted as an efficient alternative to SBS and other bootstrap approaches.

Here, we present UFBoot2, which substantially accelerates UFBoot and reduces the risk of overestimating branch supports due to polytomies or severe model violations. Additionally, UFBoot2 provides suitable bootstrap resampling strategies for phylogenomic data. UFBoot2 is 778 and 8.4 times (median) faster than SBS and RAxML rapid bootstrap on tested datasets, respectively. UFBoot2 is implemented in the IQ-TREE software package version 1.6 and freely available at http://www.iqtree.org.

---

Standard nonparametric bootstrap (SBS) (Efron 1979; Felsenstein 1985) is widely used in maximum likelihood (ML) phylogenetic analyses to estimate branch supports of a phylogenetic tree inferred from a multiple sequence alignment (MSA). To achieve this, SBS generates a large number of resampled MSAs and reconstructs an ML-tree for each bootstrapped MSA. The resulting bootstrap ML trees are then used either to compute branch supports for the ML-tree reconstructed from the original MSA or to build a consensus tree with support values.

Although fast ML-tree search algorithms exist for large datasets (Vinh and von Haeseler 2004; Stamatakis 2006; Guindon et al. 2010; Nguyen et al. 2015) SBS is still very computationally intensive. To improve computing time rapid bootstrap (RBS; Stamatakis et al. 2008) and the ultrafast bootstrap (UFBoot; Minh et al. 2013) were developed. While RBS resembles the conservative behavior of SBS (i.e., underestimating branch supports), UFBoot provides relatively unbiased bootstrap estimates under mild model misspecifications.

The key idea behind UFBoot is to keep trees encountered during the ML-tree search for the original MSA and to use them to evaluate the quality (likelihood) of the bootstrap MSAs. To speed up likelihood computation even further for bootstrap MSAs, IQ-TREE employed the resampling estimated log-likelihood (RELL) strategy (Kishino et al. 1990). For each bootstrap MSA the tree with the highest RELL score (RELL-tree) represents the ML-bootstrap tree. Contrary to SBS, UFBoot does not further ML optimize this tree. The discrepancy in branch supports between UFBoot and SBS emerges as bootstrap trees inferred by UFBoot and SBS might be different.

Here, we present UFBoot2 that substantially speeds up UFBoot and reduces the risk for overestimated branch support due to polytomies or severe model violations. We will also discuss several resampling strategies for phylogenomic data recently implemented in UFBoot2.

## Accelerating UFBoot

The likelihood computation is the major runtime bottleneck of all ML software because it lies at the core of all analyses. The pruning algorithm (Felsenstein 1981) efficiently computes the likelihood of phylogenetic trees, but still does not scale well for large data sets. Therefore, we adopted a modification to Felsenstein’s algorithm, first introduced in RAxML. The modification exploits the reversible property of models of sequence evolution typically used in phylogenetic analysis, which led to a theoretical speedup of 4 (for DNA) or 20 (for protein data) when estimating branch lengths. Moreover, we employed the SIMD (single instruction, multiple data) feature to concurrently compute the likelihood of two MSA-sites with streaming SIMD extensions or four MSA-sites with advanced vector extensions, thus leading to a theoretical speedup of two or four compared with a non-SIMD implementation. IQ-TREE code was further optimized to avoid redundant computations.

Our benchmark on 70 DNA and 45 protein MSAs showed that UFBoot2 achieved a median speedup of 2.4 times (maximum: 77.3) compared with UFBoot version 0.9.6 (released on October 20, 2013).

## Correction for polytomies

Polytomies refer to multifurcating nodes in the tree that cannot be resolved due to low phylogenetic signal in the data. However, phylogenetic reconstruction always assumes strictly bifurcating trees. When resolving polytomies there might be multiple equivalently optimal bifurcating trees. As UFBoot saves only a single optimal bifurcating tree for each bootstrap MSA, it might cause over-optimistic bootstrap supports for short branches (Simmons and Norton 2014).

To correct for this shortcoming UFBoot2 implemented the following technique. Instead of assigning the bootstrap tree with the highest RELL for each bootstrap MSA, UFBoot2 will randomly select trees encountered during tree search, whose RELL scores are less than *ε_boot_* (default: 0.5) away from the highest RELL.

It was shown with a star tree simulation (Simmons and Norton 2014) that SBS and RBS sometimes led to false positives (bootstrap supports ≥ 95% for non-existing branches), whereas with this technique UFBoot2 never supported such branches (support values ≤ 88%). Therefore, the above technique prevents over-optimistic supports.

## Reducing the impact of model violations

Minh et al. (2013) showed that severe model violations inflate UFBoot support values. To resolve this issue UFBoot2 provides an option to conduct an additional step once the tree search on the original MSA is completed. Here, the best RELL-trees are further optimized using a hill-climbing nearest-neighbor interchange (NNI) search based directly on the corresponding bootstrap MSA. Thus, this extra step operates like SBS, but with a quick tree search to save time. Bootstrap supports are then summarized from the resulting corrected bootstrap trees. In the following, we called this UFBoot2+NNI.

We repeated the PANDIT simulations in (Minh et al. 2013) to compare the accuracy of UFBoot2 and UFBoot2+NNI with SBS (1000 replicates using IQ-TREE) and RBS (RAxML bootstopping criterion). Figure 1 shows the results. If the sequence evolution model used to infer the ML-tree agrees with the model used for simulations, then SBS, RBS and UFBoot2+NNI underestimated branch supports, the latter to a lower degree (Figure 1A; curves above the diagonal), whereas UFBoot2 obtained almost unbiased branch supports (Figure 1A; curve close to the diagonal). Severe model violations do not influence SBS (Figure 1B; RBS not shown because RAxML does not support simpler model). However, UFBoot2 (like UFBoot) overestimated the branch supports (Figure 1B; curve below the diagonal), while UFBoot2+NNI only slightly underestimated the bootstrap values (Figure 1B; curve closest to the diagonal).

**Figure 1.**
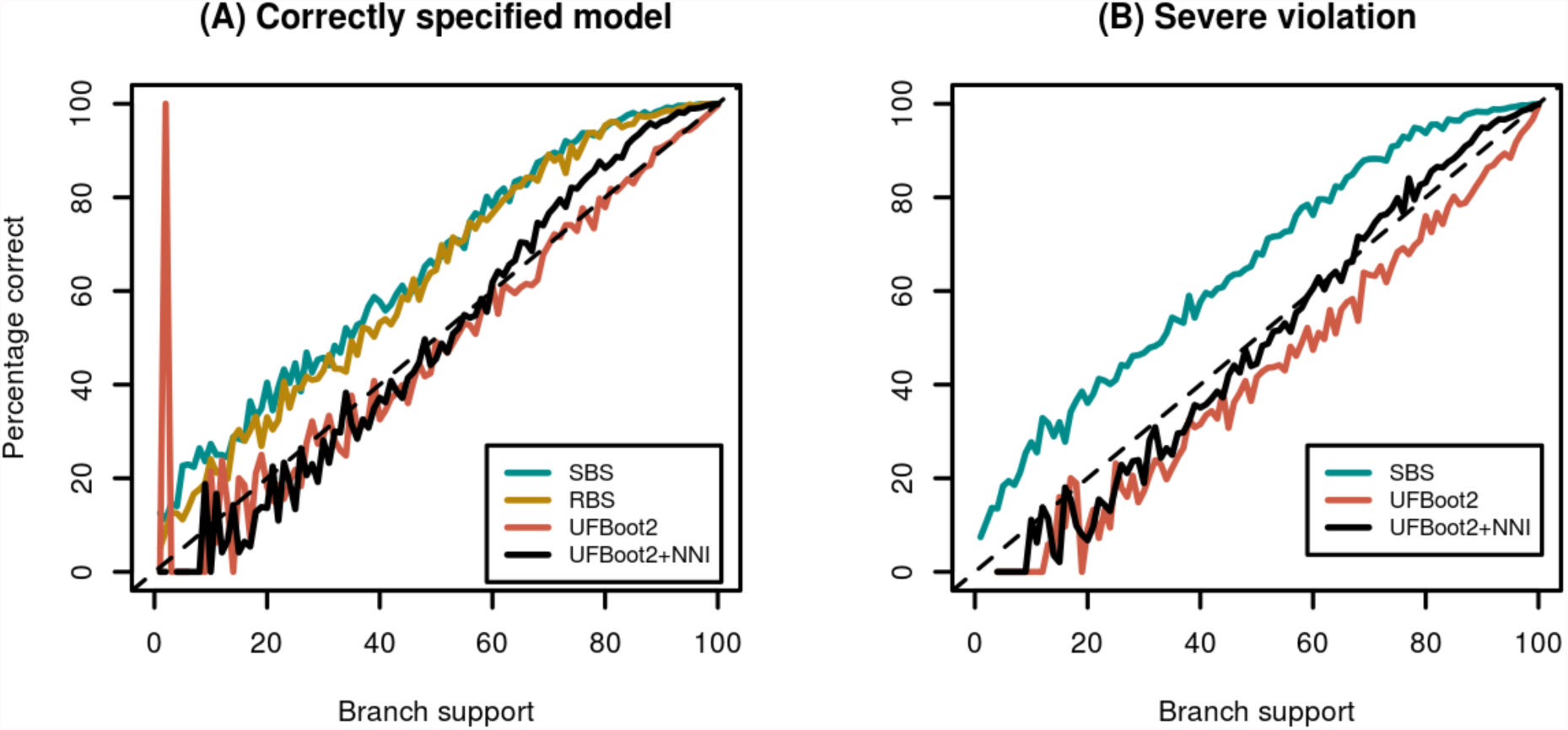
Accuracy of the standard bootstrap (SBS), RAxML rapid bootstrap (RBS), ultrafast bootstrap (UFBoot2) and UFBoot2+NNI for (A) correctly specified models and (B) severely misspecified models. Curves above the diagonal indicate underestimation of branch supports whereas curves below the diagonal indicate overestimation.

In terms of computation times, UFBoot2 and UFBoot2+NNI showed a median speedup of 778 (range: 200-1,848) and 424 (range: 233-749) compared with SBS, respectively. Compared with RBS, UFBoot2 and UFBoot2+NNI are 8.4 (range: 1.5-51.2) and 5.0 (range: 0.8-32.6) times faster, respectively. Therefore, UFBoot2+NNI is two times (median) slower than UFBoot2.

We conclude that UFBoot2 and UFBoot2+NNI are fast alternatives to other bootstrap approaches. Users are advised to apply model violation detection methods (e.g., Goldman 1993; Weiss and von Haeseler 2003; Nguyen et al. 2011) before bootstrap analyses. UFBoot2+NNI should be applied if model violations are present in the data set at hand.

## Resampling strategies for phylogenomic data

Recent phylogenetic analyses are typically based on multiple genes to infer the species tree, the so-called phylogenomics. To facilitate phylogenomic analysis, UFBoot2 implements three bootstrap resampling strategies: (i) resampling MSA-sites within partitions (denoted as MSA-site resampling), (ii) resampling genes instead of MSA-sites (gene-resampling) and (iii) resampling genes and subsequently resamples MSA-sites within each gene (gene-site resampling) (Gadagkar et al. 2005). Strategy (i) preserves the number of MSA-sites for all genes in the bootstrap MSAs, whereas strategies (ii) and (iii) will lead to different number of sites in the bootstrap MSAs.

To investigate the impact of the three resampling strategies, we reanalyzed the metazoan data with 21 species, 225 genes and a total of 171,077 amino-acid sites (Salichos and Rokas 2013). Figure 2 shows the ML tree inferred under edge-unlinked partition model (Chernomor et al. 2016), which allows separate sets of branch lengths across partitions. The tree replicates previous results (Salichos and Rokas 2013) and shows the Protostomia clade (Telford et al. 2015). However, discrepancies between resampling strategies are observed: while MSA-site and gene-resamplings obtained high supports (>95%) for branches along the backbone of the tree (Figure 2; bold lines), lower supports (80%) were estimated by gene-site resampling.

**Figure 2.**
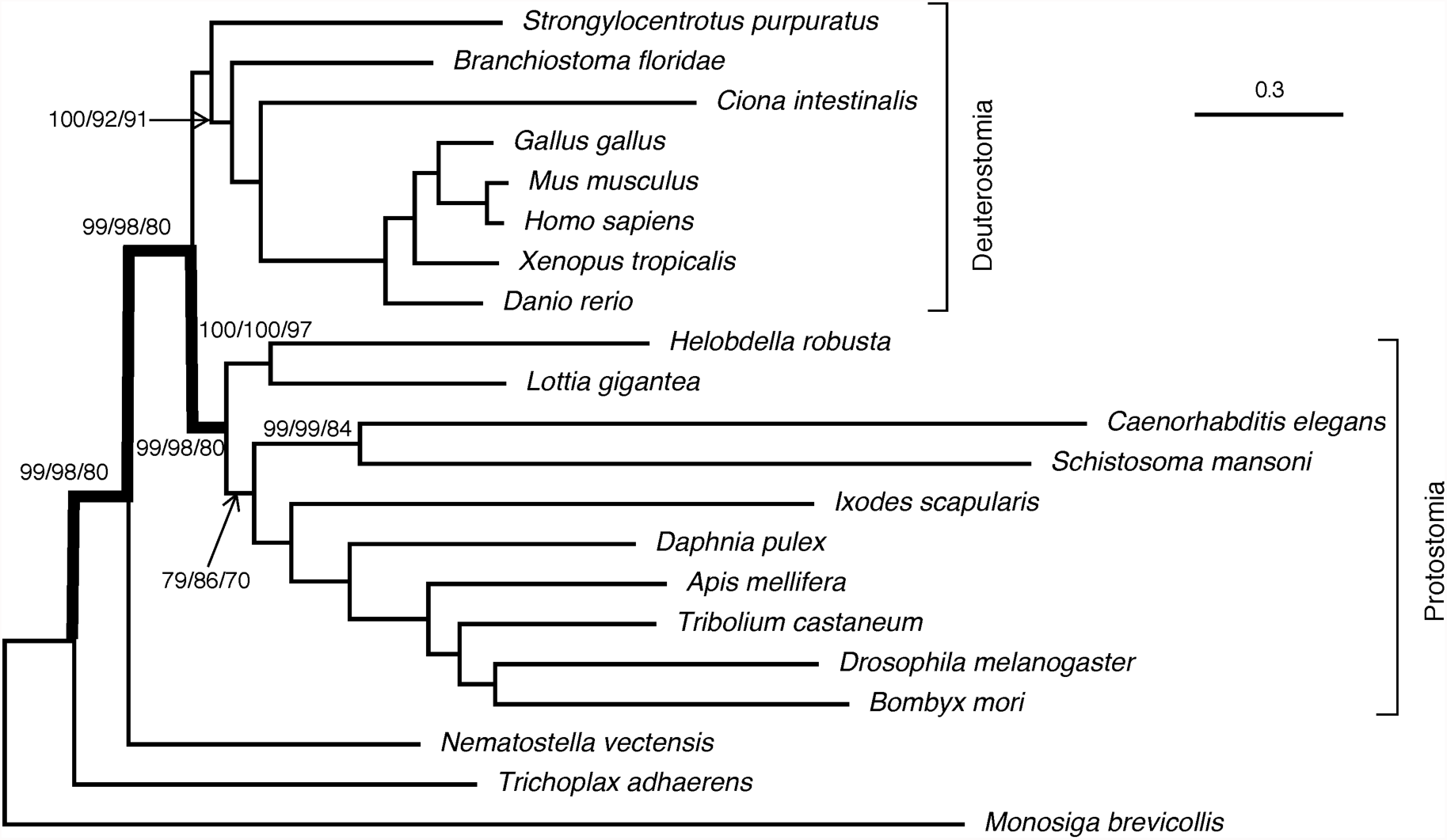
Maximum likelihood tree inferred under the edge-unlinked partition model. Numbers attached to the branches show the UFBoot2 bootstrap supports using MSA-site, gene, and gene MSA-site resampling strategies (omitted when all three supports are 100%).

By further examining 14 other empirical data sets (Bouchenak-Khelladi et al. 2008; Fabre et al. 2009; van der Linde et al. 2010; Stamatakis and Alachiotis 2010; Pyron et al. 2011; Nyakatura and Bininda-Emonds 2012; Springer et al. 2012; Hinchliff and Roalson 2013; Salichos and Rokas 2013; Dell’Ampio et al. 2014), we observed more discrepancies between resampling strategies (data not shown). Exceptionally for some data sets a number of branches showed almost no support (≤10%) for one resampling but high supports (≥95%) for the other two resampling strategies. However, there is no tendency towards systematically lower supports obtained by one resampling strategy.

Taking into account the above findings, we recommend to apply all alternative resampling strategies. If similar bootstrap supports are obtained, then one can be more confident about the results.

## Conclusions

UFBoot2 significantly improves speed and accuracy of bootstrap values compared to UFBoot. It also offers new functionalities in the presence of model violations and in its applicability to phylogenomic data. In general, since SBS, RBS and UFBoot2+NNI share a disadvantage of being conservative, more research is necessary to understand the different biases introduced by the available phylogenetic bootstrap estimation methods.

## Acknowledgment

The authors thank Stephen Crotty for comments on the manuscript. This work was supported by Vietnam National Foundation for Science and Technology Development (102.01- 2013.04). AVH, BQM, and OC were supported by the Austrian Science Fund - FWF (grant no. I-2805-B29 and I-1824-B22).

